# LizardLens: A Two-Stage Deep Learning Pipeline for Detecting and Classifying Similar Species in Visually Complex Environments

**DOI:** 10.64898/2026.06.10.731342

**Authors:** Wen Han Chia, Ilia Jahanshahi, Le Yang Loh, Anqi Zheng, Niral Verma, Stephen Mussmann, Breanna Shi, James T. Stroud

**Affiliations:** College of Computing, Georgia Institute of Technology, Atlanta GA, United States; Department of Computer Science, Iowa State University, Ames IA, United States; School of Biological Sciences, Georgia Institute of Technology, Atlanta GA, United States

**Keywords:** community science, deep learning, automated species identification, object detection, *Anolis* lizards, image classification

## Abstract

Community science platforms like iNaturalist generate unprecedented volumes of biodiversity data, but their scientific utility depends critically on accurate species identification—a persistent challenge when contributors often lack taxonomic expertise. We developed “*LizardLens*”, a two-stage machine learning pipeline that decouples object detection from species classification to enable fine-grained identification of morphologically similar organisms in visually complex field photographs. Using 10,000 verified iNaturalist images of five *Anolis* lizard species in Florida, we trained specialized YOLO-based detection and Swin Transformer classification models and compared performance against state-of-the-art single-stage architectures. Our two-stage pipeline achieved 83.0% Top-1 accuracy and a macro-averaged F1-score of 89.0%, indicating strong precision-recall performance across species and outperforming single-stage YOLOv8 and YOLOv12 models across all evaluation metrics for all species, with relative improvements ranging from 10.5% to 13.2%. Gradient-weighted Class Activation Mapping (Grad-CAM) indicated that the model’s predictions were consistently associated with regions corresponding to diagnostic morphological (e.g., head shape, feet, and limb lengths) and pattern features (e.g., ocular rings and body patterning), providing evidence that LizardLens leverages biologically relevant visual cues consistent with those used by expert taxonomists. Error analysis identified partial occlusion and multiple proximate individuals as primary sources of missed detections, while spurious detections of lizard-like environmental features (e.g., sticks, bark) represented the dominant false positive error mode. We deployed LizardLens as an accessible web application featuring interactive bounding box correction, ranked species predictions with confidence scores, directly supporting the “*Lizards on the Loose*” middle school community science initiative. By combining technical advances in fine-grained visual classification with user-centered design, LizardLens demonstrates how machine learning can simultaneously enhance data quality for biodiversity monitoring and provide authentic scientific experiences for student participants. Our approach is generalizable to other small-bodied organisms in complex habitats and provides a framework for translating computer vision advances into practical tools for community science and conservation.

## 1. Introduction

Digital imagery has revolutionized biology, enabling unprecedented scales of biodiversity data collection through community science initiatives (Chandler *et al*., 2017; Di Cecco *et al*., 2021). Platforms such as iNaturalist harness the collective power of thousands of contributors, generating organismal observation records that far exceed the capacity of professional scientists alone (Mason *et al*., 2025). These crowdsourced datasets have become invaluable resources for tracking species distributions, monitoring invasive species, and documenting phenological shifts (Encarnação, Teodósio and Morais, 2021; Jaskuła *et al*., 2021; Binley *et al*., 2023). However, the scientific utility of community science data depends critically on accurate species identification—a challenge that remains substantial when contributors lack taxonomic expertise (Koo *et al*., 2022).

Species misidentification represents a persistent bottleneck in community science platforms such as iNaturalist. While iNaturalist employs a verification system where community members review and correct uploaded identifications, this process is labor-intensive and prone to error propagation (White *et al*., 2023; Mesaglio *et al*., 2025). The problem is particularly acute for taxonomic groups with subtle morphological differences or when photograph quality is suboptimal (Jäckel *et al*., 2023; Coca-de-la-Iglesia *et al*., 2024). As community science initiatives continue to expand, developing automated tools to improve identification accuracy has become increasingly important for maintaining data quality and scientific rigor (Van Horn *et al*., 2018).

Recent advances in deep learning offer promising solutions for automated species identification from digital images. Object detection models, particularly the YOLO (“You Only Look Once”) architecture (Redmon *et al*., 2016), have demonstrated remarkable performance in rapidly localizing objects within images (Murat and Kiran, 2025). Similarly, classification algorithms employing approaches such as convolutional neural networks have achieved high accuracy in distinguishing between visually similar categories (Krizhevsky, Sutskever and Hinton, 2012). However, applying these methods to wildlife imagery presents unique challenges. Many organisms appear small relative to overall image dimensions, and natural habitats introduce substantial visual complexity through varied colors, textures, and structures (Mulero-Pázmány *et al*., 2025a). Furthermore, closely related species often exhibit superficial similarity that confounds even state-of-the-art classification models (Shen *et al*., 2025). While end-to-end models can perform classification from the full image, they often struggle with fine-grained visual discrimination when targets are small and backgrounds are complex (Mulero-Pázmány *et al*., 2025b). Decoupling these tasks into a two-stage pipeline—where specialized detection models first localize organisms and dedicated classification models subsequently identify species from cropped images—allows each component to be independently optimized, potentially improving overall performance (Ren *et al*., 2016; Mulero-Pázmány *et al*., 2025b).

Here, we developed a machine learning pipeline specifically designed to address these dual challenges of fine-grained object detection in visually complex environments and accurate classification of morphologically similar species. Our system integrates object detection and species classification to process community science photographs of *Anolis* lizards in Florida. This study system is particularly well-suited for method development, as thousands of verified iNaturalist records exist for five common species that vary in their degree of visual similarity (Van Horn *et al*., 2018). Anoles exemplify the detection and classification difficulties outlined above: they are typically small within photographs, inhabit structurally complex vegetation, and several species share overlapping morphological features (Losos, 2009; Stroud and Losos, 2020; Stroud *et al*., 2023; Price *et al*., 2025).

Our pipeline directly supports “*Lizards on the Loose*,” a community science initiative engaging middle school students in biodiversity monitoring. While this program provides valuable scientific training and generates occurrence data for both native and invasive species, student participants often lack the taxonomic expertise for confident species-level identification (Hooykaas *et al*., 2019). By providing immediate, probability-based species classifications, our LizardLens pipeline dramatically improves data quality while maintaining student engagement. This approach addresses two critical conservation objectives: first, it creates authentic research experiences that foster scientific literacy in young students (Queiruga-Dios *et al*., 2020); second, it generates high-quality verified records for four invasive anole species, supporting management efforts and invasion biology research (Encarnação, Teodósio and Morais, 2021). This integration of technological innovation with community science demonstrates how machine learning can enhance both the scientific value and educational impact of public participation in biodiversity research.

Our study had four primary objectives: (1) develop and validate a two-stage machine learning pipeline optimized for fine-grained detection and species-level classification of small animals (here, *Anolis* lizards) in visually complex field photographs; (2) compare our pipeline’s performance against state-of-the-art single-stage detection models to assess the advantages of task decoupling; (3) deploy an accessible web-based application (LizardLens) for community science use; and (4) interpret the classification model’s decision-making process through gradient-weighted class activation mapping (Grad-CAM) (Selvaraju *et al*., 2017) to understand which morphological features drive species predictions. In the following four sections, we describe our contributions towards each of these four objectives.

## 2. Methods

Our classification pipeline consisted of two sequential stages: (1) object detection to localize lizards within digital images, and (2) species-level classification of detected individuals. Below, we describe dataset assembly, model architecture, training procedures, and performance evaluation for each stage.

### 2.1. Training dataset assembly

We assembled a training dataset from iNaturalist photographic records of the five most common *Anolis* lizard (anole) species in Florida: the American green anole (*Anolis carolinensis*), Puerto Rican crested anole (*A. cristatellus*), Hispaniolan bark anole (*A. distichus*), Cuban knight anole (*A. equestris*), and Cuban brown anole (*A. sagrei*). For each species, we filtered for “Research Grade” images, photographs that have received community verification for correct species identification, and downloaded the images from the iNaturalist records using the API link (*Export Observations, iNaturalist*, 2026). As of 17th Jan 2025, the image counts of brown anole, green anole, crested anole, bark anole and knight anole are 44756, 44242, 6075, 2671 and 2301 respectively. Next, most of the anoles‘ bounding boxes were automatically annotated using GroundingDINO (Liu *et al*., 2024), a zero-shot Vision-Language model on Roboflow (Dwyer, Nelson and Hansen, 2026). The remaining unannotated anoles images were manually annotated by the authors (W.H.C., A.Z., N.V.). Next, the authors then retained only images that display distinct species features as stated in the Florida Anole field identification guide on iNaturalist (Thawley and Stroud, 2026). To mitigate the issue of class imbalance, we included only 2000 images of each species, yielding a total dataset count of 10,000 images.

Images were randomly partitioned into training (n = 8,000; 80% of photos), validation (n = 1,000; 10%), and test (n = 1,000; 10%) subsets, stratified by species to maintain equal class representation (1,600 training, 200 validation, and 200 test images per species). Given that some images contain more than one anole lizard detection, the per-species lizard detection counts slightly exceed 2000 (Table 1). However, the resulting class imbalance is negligible (maximum deviation <1% from the mean). Note that all of the images with multiple anole lizard detections contain only one species; no images contain more than one anole species.

**Table 1.** Dataset composition by species. Images were curated to 2,000 per species and partitioned into training (80%; n = 1600), validation (10%; n = 200), and test (10%; n = 200) image subsets. The number of lizards detected (Lizards detected Total) exceeds the number of images analyzed (Images Total) because some images contain more than one anole in them (all images contain at least one). Training, Validation, and Test columns report lizard detection counts per split.

| Dataset | Images<br>(Total) | Lizards detected<br>(Total) | Lizards detected<br>in Training images | Lizards detected<br>in Validation<br>images | Lizards detected<br>in Test images |
| --- | --- | --- | --- | --- | --- |
| Bark Anole | 2000 | 2018 | 1614 | 202 | 202 |
| Brown Anole | 2000 | 2009 | 1606 | 203 | 200 |
| Crested Anole | 2000 | 2018 | 1609 | 205 | 204 |
| Green Anole | 2000 | 2041 | 1636 | 202 | 203 |
| Knight Anole | 2000 | 2028 | 1625 | 201 | 202 |
| Total | 10000 | 10114 | 8090 | 1013 | 1011 |

### 2.2. Stage 1: Detection model training

The primary objective of the object detection stage was to localize lizards within input images and generate bounding boxes around detected individuals. We evaluated three state-of-the-art object detection architectures: YOLOv8x with 68.2 million parameters (Varghese and M., 2024), YOLOv11x with 56.9 million parameters (Khanam and Hussain, 2024), and Real-Time Detection Transformer RT-DETR(L) with 42 million parameters (Zhao *et al*., 2024). The best-performing model was selected for integration into the classification pipeline. For object detection training, we created a modified version of the dataset by replacing species-level labels with a single “lizard” class label, enabling the model to detect Anolis lizards regardless of species identity. Each model was fine-tuned with the training dataset for 100 epochs using a batch size of 128 and Stochastic Gradient Descent (SGD) as the optimizer. Input images were resized to 640x640 using letterbox padding to preserve aspect ratio. Model training was conducted on two A100 GPUs with 80GB VRAM using Ultralytics package (Jocher, Qiu and Chaurasia, 2023).

### 2.3. Detection model performance evaluation

We evaluated detection performance using precision, recall, F1-score at an evaluation IoU threshold of 0.5, and mean average precision (mAP) calculated across intersection over union (IoU) thresholds ranging from 0.50 to 0.95 (mAP@50-95) (Padilla *et al*., 2021) (Equations 1 - 4). During inference, non-maximum suppression (NMS) was performed at an IoU threshold of 0.7 to remove redundant overlapping detections prior to evaluation. Precision quantifies the proportion of correct detections among all predicted bounding boxes, while recall measures the proportion of actual lizards successfully detected, revealing whether the model systematically misses certain species (Sokolova and Lapalme, 2009). F1-score provides a harmonic mean of precision and recall, and mAP@50-95 integrates precision across multiple IoU thresholds to assess both localization accuracy and detection sensitivity (Powers, 2020; Padilla *et al*., 2021).

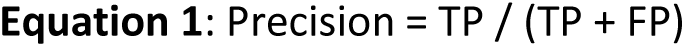

where TP is true positives (correct predictions for the detections) and FP is false positives (incorrect predictions assigned to the detections). Recall measures the proportion of actual lizards of each species that were correctly identified, revealing whether the model systematically misses certain species:

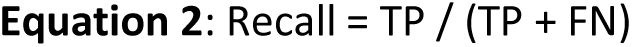

where FN is false negatives (lizard detections incorrectly assigned to other classes in test images). F1-score provides a harmonic mean of precision and recall, offering a balanced assessment of model performance that accounts for both prediction accuracy and detection sensitivity:

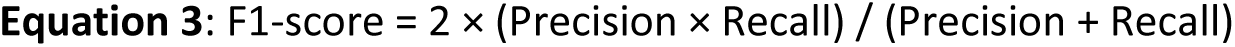

Mean average precision (mAP) is computed by first calculating the average precision (AP) for each class as the area under its precision–recall curve, then averaging AP values across all classes. mAP@50–95 averages these values across ten evaluation IoU thresholds from 0.50 to 0.95 (in 0.05 steps), providing a stricter measure of localization quality than single-threshold mAP (Padilla et al., 2021):

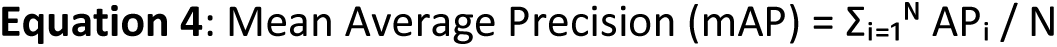

where N is the number of classes and APᵢ is the average precision for class i.

We prioritized two metrics when selecting the optimal detection model. First, recall was considered paramount because missed detections in this initial stage prevent any downstream classification, causing complete pipeline failure for those images. High recall ensures maximum retention of lizard-containing images for subsequent species classification. Second, mAP@50-95 was emphasized because accurate bounding box localization is critical for preserving diagnostic morphological features in cropped images passed to the classification model. Tight bounding boxes minimize extraneous background information that could confound species classification, while loose boxes risk excluding key identifying characteristics.

The following metrics were recorded: precisions of 0.946, 0.933, 0.965, recall of 0.942, 0.932, 0.938, F1-score of 0.944, 0.935, 0.951 and mAP@50-95 of 0.808, 0.800, 0.796 for YOLOv8x, YOLOv11x and RT-DETR(L) respectively. As such, the YOLOv8x model was selected as the first stage of the pipeline.

### 2.4. Stage 2: Classification model training and evaluation

Following object detection, cropped lizard images were passed to an anole species classification model to assign probability-based identifications to one of five Anolis species. We fine-tuned and evaluated the performances of 3 classification models, data-efficient image transformers (DeiT) base model with 87 million parameters (Touvron *et al*., 2021), Vision Transformer (ViT) base model with 86.9 million parameters (Dosovitskiy *et al*., 2021) using pre-trained weights from the google/vit-base-patch16-224 checkpoint available on Hugging Face (Deng *et al*., 2009; Wu *et al*., 2020), and Swin Transformer base model with 88 million parameters (Liu *et al*., 2021). For classification model fine-tuning, we created a modified version of the dataset by cropping images of the anoles based on the bounding box annotations in the original dataset. The original species labels were retained. Each model was trained for 30 epochs using Adam optimization, image size of 384, batch size of 128, and HuggingFace package (Wolf *et al*., 2020). Two H100 GPU with 80GB VRAM were utilized to support this operation.

We evaluated classification performance using top-1 accuracy (Equation 5) together with per-species precision, recall, and F1-score. Top-1 accuracy measures the proportion of test images for which the model’s highest-confidence class prediction matches the ground-truth label, providing an overall measure of classification performance. Per-species precision was emphasized as an indicator of prediction reliability for each species.

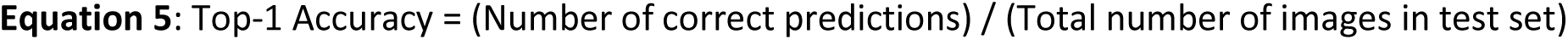

where a top-1 prediction is the class assigned the highest confidence score by the model, and a prediction is counted as correct if it matches the ground-truth label of the anole instance.

The performance of the models, DeiT, ViT and Swin Transformer were logged respectively: top-1 accuracy of 0.907, 0.919 and 0.939. Furthermore, the precision, recall and F1-score of each class were logged as shown in Table 2. Swin Transformer scored a high precision (above 90%) for almost all classes except for Brown anole. Coupled with a high top-1 accuracy of 0.939, the Swin Transformer was selected to be the classification model for the pipeline.

**Table 2:**
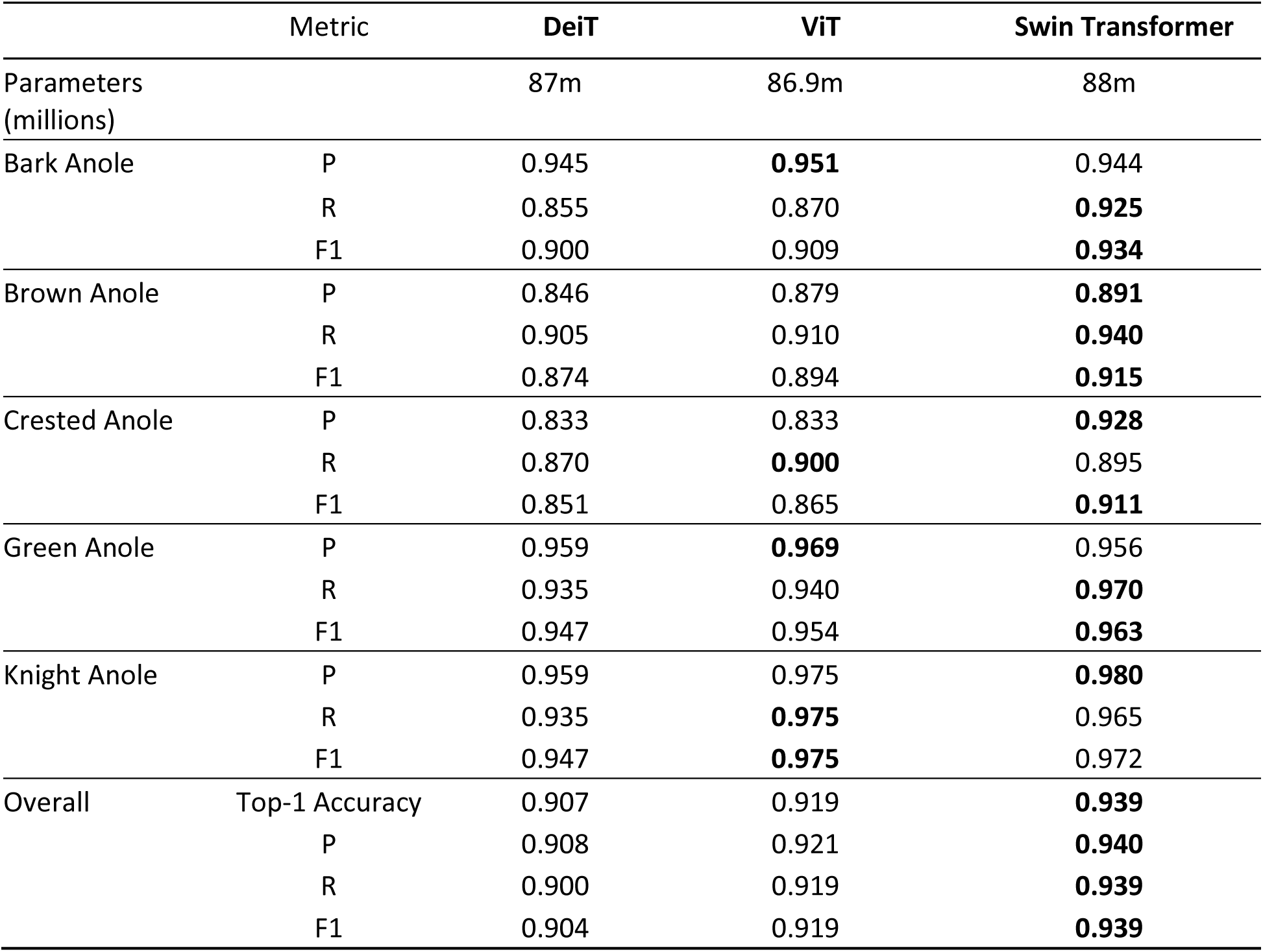
The performance of each classification model fine-tuned with a dataset consisting of cropped anole lizards detected in test images. The precision, recall and F1-score of each anole class were reported for each model.

### 2.5. Final Model Performance Evaluation

We wanted to compare the performance of LizardLens, the two-stage classification pipeline consist of fine-tuned YOLOv8x detection model and Swin Transformer classification model, against other single-stage detection model. YOLOv8x serves a dual role in our evaluation: it is the detection backbone within our two-stage pipeline, and it is evaluated independently as a single-stage baseline. This controlled comparison isolates the contribution of the dedicated classification stage — since the same detector produces the bounding boxes in both settings, any performance difference is directly attributable to replacing YOLOv8x’s built-in classification head with the Swin Transformer backbone. This ablation-style design answers a core question of this study: whether the added complexity of a two-stage pipeline yields meaningful gains in fine-grained species discrimination over a single-stage approach. Secondly, YOLOv12 introduces an attention-centric architecture that departs from the purely convolutional feature extraction used in prior YOLO variants (Tian, Ye and Doermann, 2025). Because attention mechanisms can capture long-range spatial dependencies between discriminative features — such as dewlap coloration, dorsal patterning, and head scale morphology — this architecture is a compelling baseline for fine-grained species classification tasks where subtle inter-class differences are spatially distributed across the image. As such, the YOLOv12x and YOLOv8x models were chosen as the single-stage detection models of comparison against the two-stage classification pipeline, LizardLens. They were fine-tuned with the original training dataset (n = 8090) and their performance was evaluated on the held-out test set using standard classification metrics such as top-1 accuracy and macro-averaged precision, recall, and F1-score (Sokolova and Lapalme, 2009). Top-1 accuracy measures the proportion of images for which the model’s highest-probability prediction matched the ground truth species label at the detection level. The evaluation results are shown in Section 3.1.

### 2.6. LizardLens: Pipeline Deployment and Accessibility

To make our validated classification pipeline accessible to community scientists and educators, we developed a web-based application for LizardLens that processes field photographs and returns immediate species identifications. The application was designed specifically to support non-expert users—including middle school students participating in the Lizards on the Loose initiative—by providing probability-based species classifications that enhance identification accuracy while maintaining user engagement. The complete user workflow is illustrated in Figure 1.

**Figure 1.**
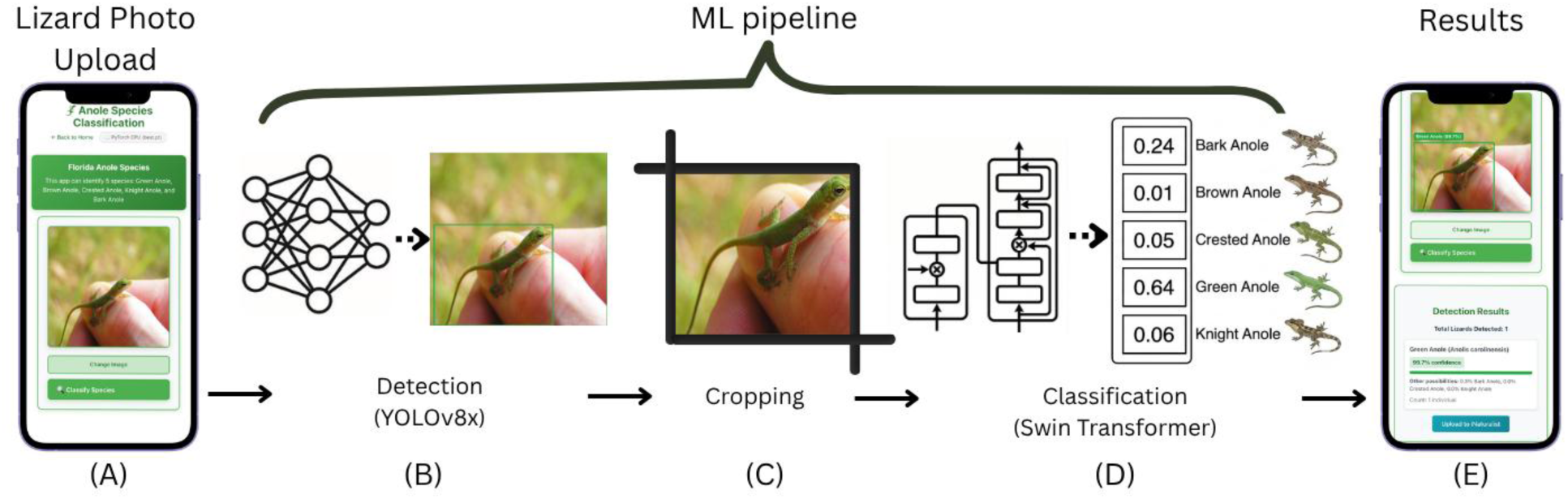
End-to-end web application workflow. (A) Users upload a field image. (B) The backend detects anoles and produces bounding boxes. (C) Cropped lizard detections are passed to the species classifier. (D) The Swin Transformer performs species classification. (E) The frontend visualizes bounding boxes and per-lizard detection confidence scores before optionally submitting validated observations to iNaturalist.

Figure 1. End-to-end web application workflow. Users upload a field image, the backend detects anoles and produces bounding boxes. Cropped lizard detections are passed to the species classifier. The frontend visualizes bounding boxes and per-lizard detection confidence scores before optionally submitting validated observations to iNaturalist.

#### 2.6.1. LizardLens: System Architecture

The application consists of a React user interface and a FastAPI (Ramírez, 2026) python server hosting the trained detection and classification models. The system is deployed on Georgia Tech’s College of Computing server and accessible at (*Lizard Lens*, 2026). When users upload photographs, images are transmitted to the server via HTTP POST, processed through the pipeline, and results are returned to the client interface in JSON format. The client application transforms the JSON response into a list of predictions. It then takes these predictions to render Bounding Boxes over the original image, color-coded by confidence level, with labels displaying the species name and confidence percentage.

#### 2.6.2. LizardLens: User Workflow

The application guides users through four sequential stages (Figure 1):

1. Image Upload and Automated Inference. Users upload field photographs through the web interface which accepts standard JPG and PNG formats. Upon submission, the back-end detection model (Stage 1) localizes individual lizards and generates bounding boxes, which are then passed to the classification model (Stage 2) for species-level predictions. Inference typically completes within 1.28 seconds per image on a standard CPU (modern Intel Core/Xeon with 4GB+ RAM).
2. Interactive Detection Review. The interface displays the original photograph with color-coded bounding boxes overlaid on each detected lizard, colored by confidence level (Green for high confidence > 80%, Yellow for medium > 60%, and Red for low ≤ 60%). Users can visually assess detection quality and manually adjust bounding boxes by resizing or repositioning to correct incomplete or inaccurate detections (e.g., boxes that exclude diagnostic features or include excessive background). When a bounding box is edited, the system automatically re-extracts the cropped region and re-runs the classification model to update the species prediction based on the corrected detection.
3. Species Identification and Confidence Assessment. For each individual lizard that LizardLens detects, the application presents: (i) the predicted species common name and scientific name (i.e., Genus, species), (ii) the model’s confidence score (ranging from 0–100%), and (iii) a ranked list of alternative species predictions with associated confidence values. This transparency allows users to evaluate prediction certainty and recognize ambiguous classifications that may warrant expert verification.
4. Data Submission and Community Impact Tracking. Validated observations are submitted directly to the iNaturalist platform. Upon successful submission, the community metrics dashboard reflects the new contribution through real-time counters and species tallies, providing users with immediate scientific validation and visual confirmation of their role within the broader research initiative.

#### 2.6.3. iNaturalist Integration and Real-Time Community Dashboard

Integration with the iNaturalist platform via its public API facilitates streamlined biodiversity data contribution and amplified community impact (Callaghan *et al*., 2022). Once a user validates an automated detection and species prediction, the application enables a direct-to-iNaturalist submission workflow. This process encapsulates the field photograph and predicted species identification into an iNaturalist observation record. The development of a live community metrics dashboard (Figure 2A) fosters participation and provides immediate feedback. Leveraging React-based animation libraries (e.g., Framer Motion and React-Countup), the dashboard retrieves real-time statistics from the iNaturalist API, including total observations, unique species counts, and contributor totals across the Florida Anolis dataset. The implementation of optimistic UI updates ensures that these counters increment instantly upon a successful upload request, providing participants with a responsive experience that bridges the gap between individual contribution and collective scientific impact (Jennett *et al*., 2016).

**Figure 2.**
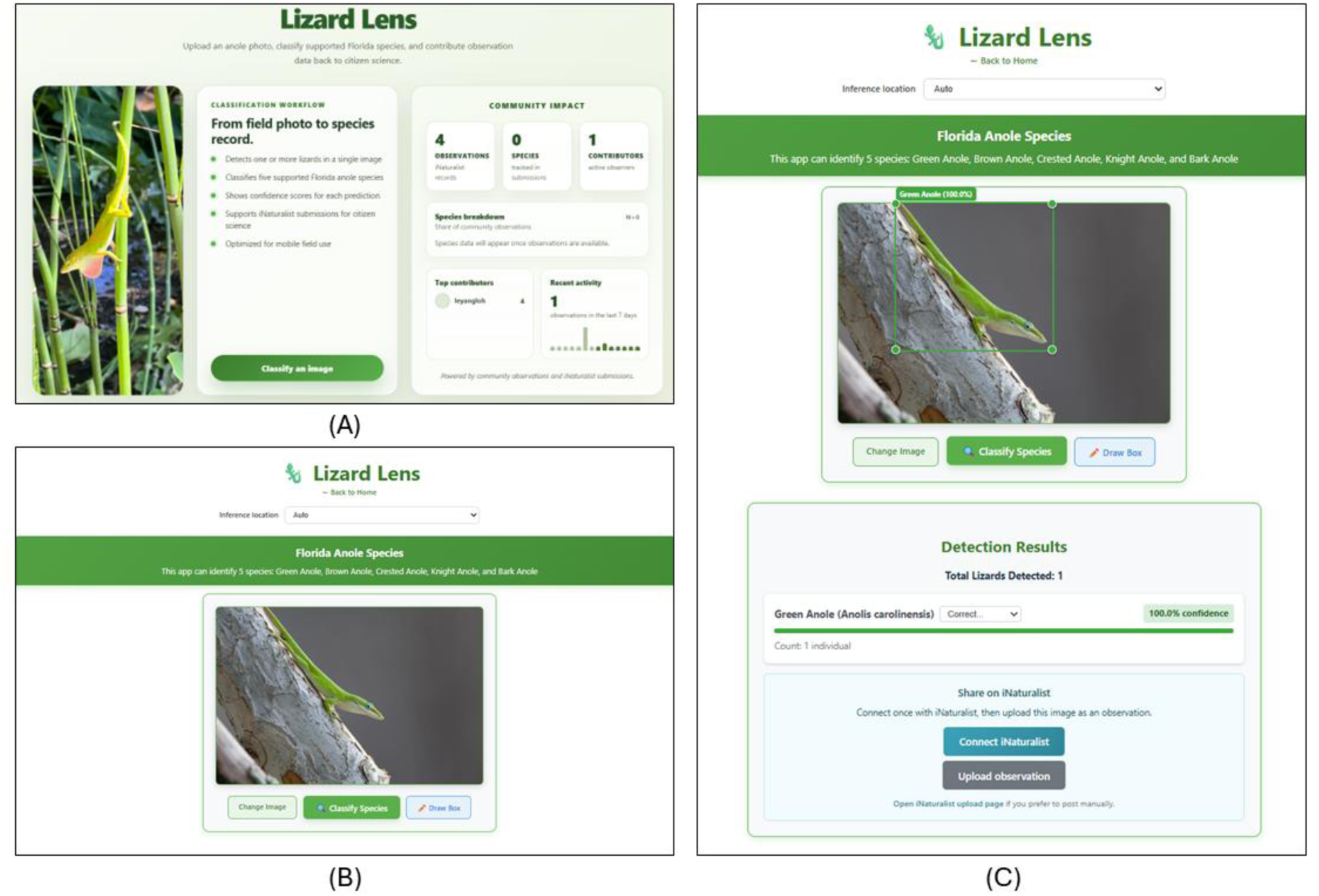
Screenshots of the LizardLens web application. (A) Homepage of the web application – a user-friendly interface that displays recent iNaturalist submissions made through the application. (B) Displays the captured image and options to change image, classify the species or manually annotate the bounding box. (C) The LizardLens prediction of the anole bounding box, confidence score, number of species and species classification are displayed. User can edit the predicted bounding box or choose to upload this observation to iNaturalist.

Figure 2. Screenshots of the LizardLens web application. (A) Homepage of the web application – a user-friendly interface that displays recent iNaturalist submissions made through the application. (B) Displays the captured image and options to change image, classify the species or manually annotate the bounding box. (C) The LizardLens prediction of the anole bounding box, confidence score, number of species and species classification are displayed. User can edit the predicted bounding box or choose to upload this observation to INaturalist.

### 2.7. Model Interpretability Analysis

To understand which morphological features the classification model relies on for species discrimination, we applied Gradient-weighted Class Activation Mapping (Grad-CAM) to visualize spatial attention patterns during inference. Grad-CAM generates heatmaps highlighting image regions that strongly influence the model’s predictions, enabling post-hoc interpretation of the classification decision process(Selvaraju *et al*., 2017; Chattopadhay *et al*., 2018).

We implemented Grad-CAM using the fine-tuned Swin Transformer as mentioned in Section 2.4. Because Swin Transformer employs a hierarchical encoder architecture incompatible with standard Grad-CAM implementations, we developed a custom reshape function to convert the model’s multi-scale feature representations into activation maps suitable for gradient-based attention visualization. We generated Grad-CAM heatmaps at multiple encoder layers and evaluated which layer produced the most biologically interpretable attention pattern. Heatmaps derived from earlier layers exhibited diffuse attention patterns dominated by low-level texture, while later layers often produced overly coarse activations. We selected the third encoder stage for detailed analysis because it consistently produced spatially localized heatmaps centered on the lizard body while preserving sufficient anatomical detail.

To assess whether the model attended to species-diagnostic morphological features, we randomly sampled 25 test images (5 per species) and J.T.S. qualitatively evaluated their corresponding Layer 3 Grad-CAM heatmaps against established identification criteria.

## 3. Results

### 3.1. LizardLens: Overall Pipeline Performance

The LizardLens two-stage pipeline outperformed both single-stage baselines across all evaluation metrics (Table 3). LizardLens achieved an F1-score of 0.890 and top-1 accuracy of 83.0%, compared with 0.802 and 74.0% for YOLOv8x and 0.724 and 68.0% for YOLOv12x. Relative improvements over the next-best model (YOLOv8x) ranged from 10.5% (precision) to 13.2% (mAP@50-95), with consistent gains across all metrics including localization accuracy (mAP@50: 0.891 vs. 0.794; mAP@50-95: 0.704 vs. 0.622).

**Table 3.** Performance comparison of the LizardLens two-stage pipeline against single-stage YOLOv8x and YOLOv12x models on the test dataset (n = 1,011, Table 1). Asterisks (*) indicate best model performance for each metric. The final column represents the relative performance improvement of LizardLens over the next-best model.

| Performance metric | YOLOv8x | YOLOv12x | **LizardLens** | LizardLens improvement |
| --- | --- | --- | --- | --- |
| Parameters | 68.2M | 59.1M | **156.2M** | - |
| Top-1 Accuracy | 0.740 | 0.680 | **0.830\*** | +12.2% |
| F1-score | 0.802 | 0.724 | **0.890\*** | +11.0% |
| Precision | 0.798 | 0.736 | **0.882\*** | +10.5% |
| Recall | 0.806 | 0.720 | **0.896\*** | +11.2% |
| mAP@50 | 0.794 | 0.676 | **0.891\*** | +12.2% |
| mAP@75 | 0.679 | 0.580 | **0.758\*** | +11.6% |
| mAP@50–95 | 0.622 | 0.533 | **0.704\*** | +13.2% |

These gains can be attributed directly to the dedicated classification stage. Because YOLOv8x serves as both the detection backbone within LizardLens and as an independent single-stage baseline, the two pipelines share identical bounding box predictions — any performance difference is therefore attributable solely to replacing YOLOv8x’s built-in classification head with the Swin Transformer. The magnitude of improvement across all metrics confirms that decoupling detection from classification and optimizing each task independently yields substantial gains in fine-grained species discrimination. Notably, YOLOv12x underperformed even the convolutional YOLOv8x despite its attention-centric architecture, which can in principle capture long-range spatial dependencies between discriminative features such as dewlap coloration and dorsal patterning (Tian, Ye and Doermann, 2025). This result suggests that architectural capacity for modeling spatial relationships does not, on its own, compensate for the lack of a dedicated classification stage when targets are small and interspecific differences are subtle.

### 3.2. Species-Level Performance

LizardLens’s performance varied among the five *Anolis* species but remained consistently high (Table 4). All species achieved precision, recall, and F1-scores exceeding 0.85, indicating robust classification regardless of species identity. In contrast, the single-stage YOLOv8x and YOLOv12x models exhibited substantially lower and more variable performance across species, with several species falling below 0.80 for multiple metrics. The pipeline performed best for *A. distichus* (F1-score = 0.930; Table 4), but was strong for all species (range = 0.850 [*A. sagrei*] - 0.920 [*A. cristatellus*]). Differences between species likely represent both biological (high intraspecific variation) and technical (bad photograph quality) features.

**Table 4.** Species-level performance comparison of the LizardLens two-stage pipeline against single-stage YOLOv8x and YOLOv12x models on the test dataset (n = 1,011; Table 1). Asterisks (*) indicate best model performance for each metric. Precision, recall, and F1-score are represented as P, R, and F1 respectively. The overall precision, recall, and F1-scores are calculated using macro-average.

| Species | Metric | YOLOv8x | YOLOv12x | LizardLens |
| --- | --- | --- | --- | --- |
| Bark Anole | P | 0.790 | 0.780 | **0.930\*** |
|  | R | 0.800 | 0.710 | **0.930\*** |
|  | F1 | 0.790 | 0.740 | **0.930\*** |
| Brown Anole | P | 0.820 | 0.730 | **0.850\*** |
|  | R | 0.740 | 0.630 | **0.850\*** |
|  | F1 | 0.780 | 0.680 | **0.850\*** |
| Crested Anole | P | 0.810 | 0.690 | **0.870\*** |
|  | R | 0.830 | 0.740 | **0.920\*** |
|  | F1 | 0.820 | 0.710 | **0.900\*** |
| Green Anole | P | 0.810 | 0.760 | **0.890\*** |
|  | R | 0.820 | 0.720 | **0.890\*** |
|  | F1 | 0.820 | 0.740 | **0.890\*** |
| Knight Anole | P | 0.760 | 0.720 | **0.870\*** |
|  | R | 0.840 | 0.800 | **0.890\*** |
|  | F1 | 0.800 | 0.750 | **0.880\*** |
| Overall | P | 0.798 | 0.736 | **0.882\*** |
|  | R | 0.806 | 0.720 | **0.896\*** |
|  | F1 | 0.802 | 0.724 | **0.890\*** |

### 3.3. LizardLens: Error Analysis

Our analysis of pipeline errors revealed specific failure modes that inform both the interpretation of performance metrics and potential avenues for improvement. We categorized errors into three primary types: missed detections (false negatives), spurious detections (false positives), and species misclassifications among correctly detected lizards.

#### 3.3.1. Overview of Error Sources

The normalized confusion matrix for the LizardLens pipeline illustrates the distribution of these error types (Figure 3). Missed detections (false negatives, Background column) occurred when the Stage 1 detection model failed to localize lizards present in images, preventing downstream classification. These errors reduced recall, F1-score, and mAP values. Spurious detections (false positives, Background row) occurred when the detection model incorrectly identified non-lizard image regions as lizards, which were subsequently passed to the classifier and assigned species labels. These errors reduced precision, F1-score, and mAP. Species misclassifications (non-diagonal elements) occurred when the Stage 2 classification model assigned incorrect species identities to correctly detected lizards. Together, these three error sources contributed to the reduction in overall Top-1 accuracy from perfect classification.

**Figure 3.**
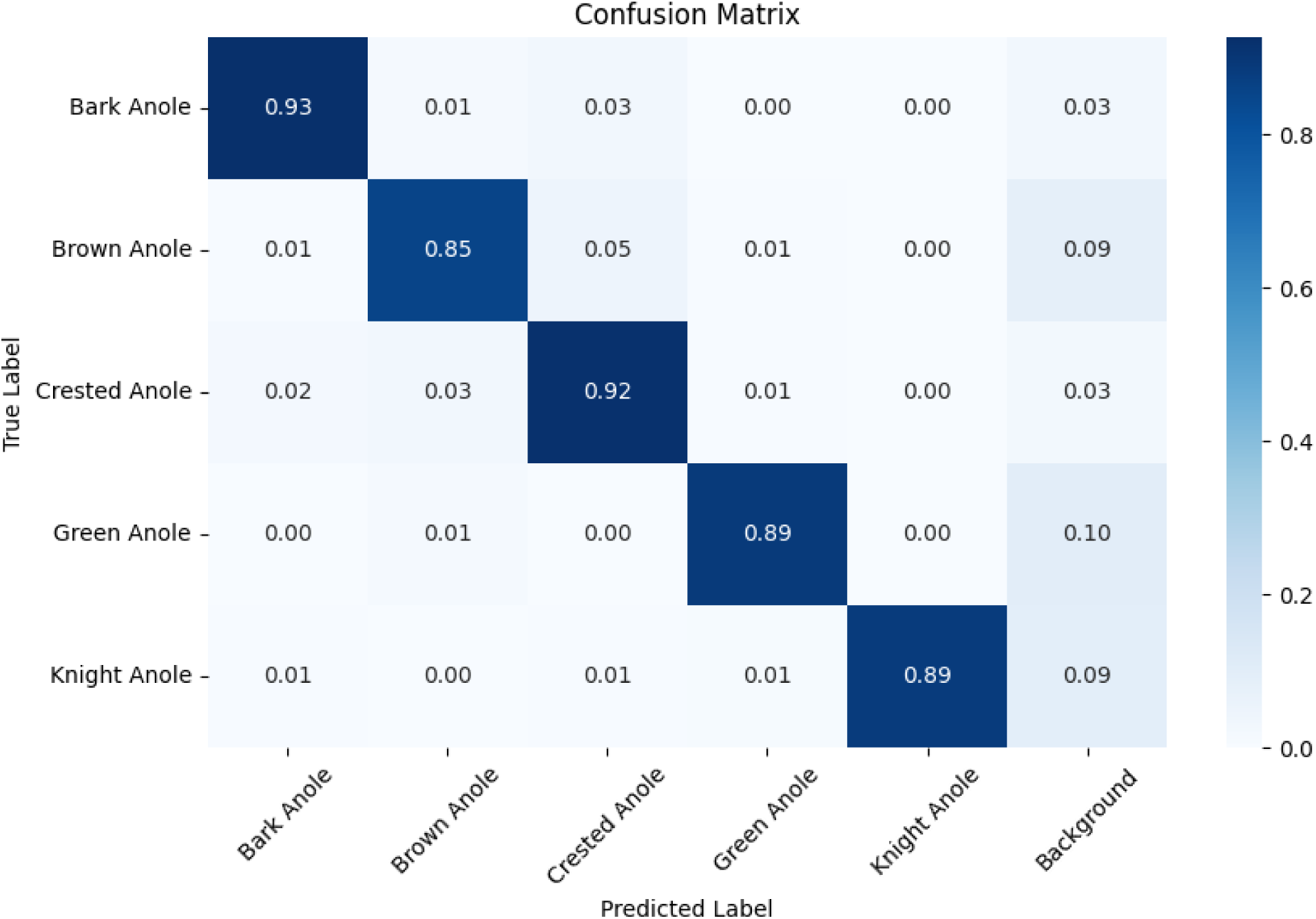
The normalized confusion matrix of the classification pipeline predictions of each anole species.

Quantitatively, out of the 1011 anole lizard detections in the test image dataset, 67 errors (6.63%) resulted from missed detections, 81 errors (8.01%) from detections, and 39 errors (3.86%) from species misclassifications among correctly detected lizards as shown in the confusion matrix (Figure 4). Among species-level classification errors, the most frequent confusions (n = 10) occurred between brown anoles (*A. sagrei*) and crested anoles (*A. cristatellus*), which exhibit exceptional similarity in body shape, coloration, dorsal patterning, and habitat use (Stroud *et al*., 2024) and can be difficult for even expert biologists to distinguish (Stroud pers. obs.). Notably, errors involving misclassification of non-lizard background objects as lizards outnumbered confusions between any pair of lizard species, indicating that distinguishing lizards from visually similar environmental features represents a greater challenge than discriminating among *Anolis* species.

**Figure 4.**
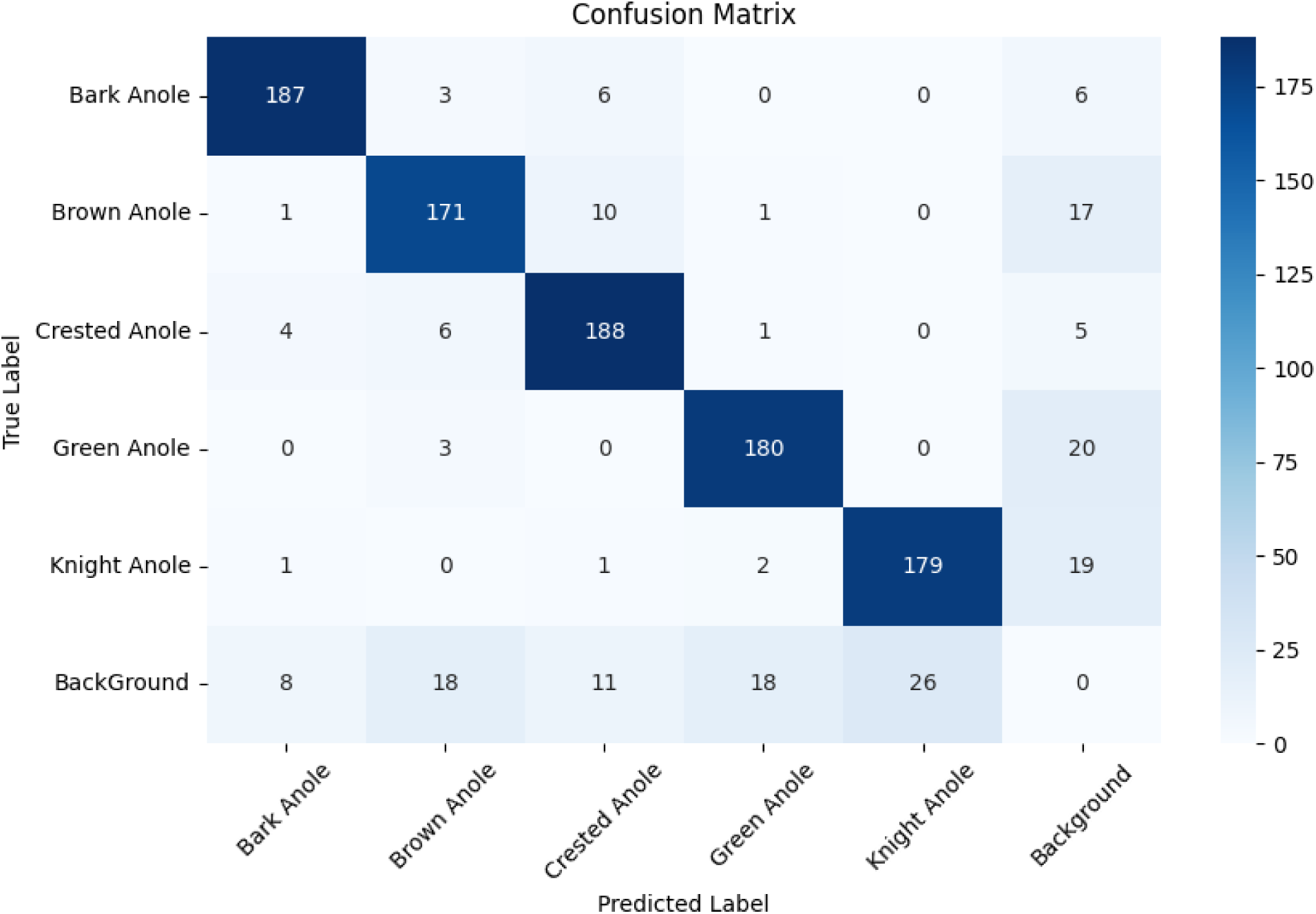
Confusion matrix, in absolute counts, of the classification pipeline’s species predictions for each anole class. Out of 1,011 anole lizard detections in the test image dataset, 81 errors (8.01%) were false positive detections, 67 errors (6.63%) were missed detections, and 39 errors (3.86%) were species misclassifications among correctly detected lizards.

Figure 3. The normalized confusion matrix of the classification pipeline predictions of each anole species.

Figure 4. Confusion matrix, in absolute counts, of the classification pipeline’s species predictions for each anole class. Out of 1,011 anole lizard detections in the test image dataset, 81 errors (8.01%) were false positive detections, 67 errors (6.63%) were missed detections, and 39 errors (3.86%) were species misclassifications among correctly detected lizards.

#### 3.3.2. False Negative Errors: Missed Detections

False negatives occurred when the Stage 1 detection model either failed to generate bounding boxes around lizards or produced boxes with insufficient overlap (IoU < 0.7) with ground truth annotations. Through qualitative analysis of error cases, we identified two primary causes:

##### Partial Occlusion

When lizards were partially obscured by vegetation, substrate, or other environmental features, the detection model frequently failed to localize individuals (Figure 5A). Images where only the head or a small body portion was visible proved particularly challenging, likely because the majority of training images depicted largely unobstructed lizards with most body regions visible.

**Figure 5.**
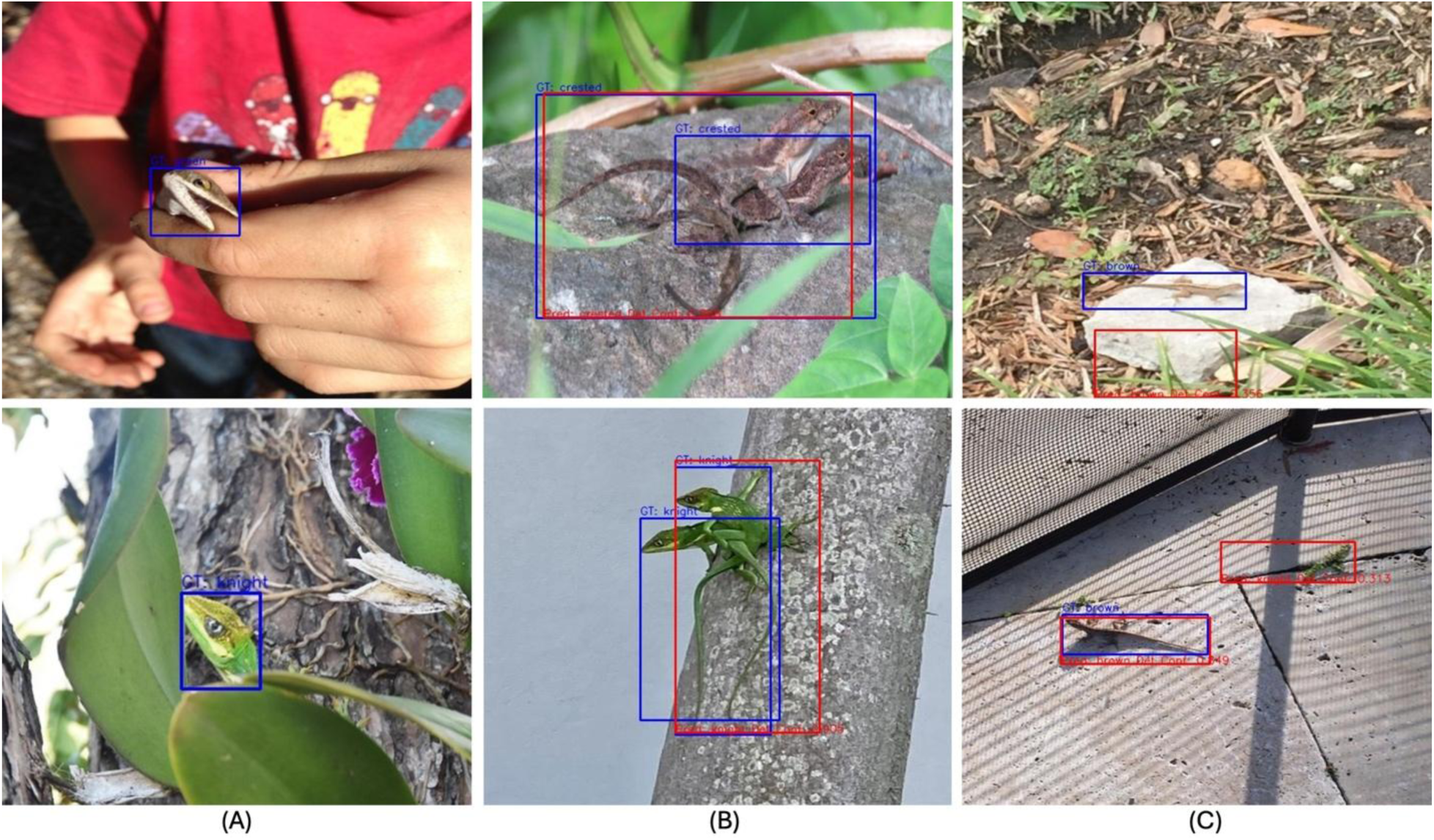
Representative error modes in the detection model. (A) False-negative due to partial occlusion (n=41 images, 4.06% of test set): the model fails to detect a lizard when only a small portion of the body (e.g., head) is visible and obscured by surrounding vegetation or substrate. (B) False-negative due to multiple proximate individuals (n=9 images, 0.89% of test set): the model generates a single bounding box encompassing two closely positioned lizards rather than detecting each individual separately. (C) False-positive detection: the model incorrectly identifies non-lizard objects (e.g., sticks, bark, or lighting patterns) as lizards, often with low confidence. Blue boxes indicate predicted detections, and red boxes indicate ground truth annotations.

##### Multiple Proximate Individuals

When multiple lizards appeared in close spatial proximity in the same photograph, the detection model often generated a single bounding box encompassing multiple individuals rather than separate boxes for each lizard (Figure 5B). This error likely stems from the Non-Maximum Suppression (NMS) post-processing step, which removes overlapping predictions above a defined IoU threshold (currently set at 0.7). When two lizards are closely positioned, their predicted bounding boxes may exceed this overlap threshold, causing NMS to retain only the highest-confidence prediction and suppress the others.

For our primary use case—species-level classification for community science—this error mode has minimal practical impact because the co-occurring lizards are conspecific. As mentioned in Section 2.1, all of the images with multiple anole lizard detections contain only the same species class. This suggests that missing one of two conspecific individuals does not substantially reduce observation quality. Importantly, most online community science initiatives, such as iNaturalist, do not allow multi-species photographic records, so photographs with more than one species are unable to be contributed.

#### 3.3.3. False Positive Errors: Spurious Detections

False positives occurred when the detection model incorrectly identified non-lizard image regions as lizards (Figure 5C), which were subsequently passed to the classification model and assigned species labels. Post-hoc analysis of confidence distributions (Figure 6) revealed that false positive predictions exhibited substantially lower and more variable detection confidence (median 0.54, IQR 0.39–0.73) compared with true positive detections, which were tightly clustered at high confidence (median 0.91, with 75% of cases exceeding 0.86). The spuriously detected objects often resembled lizards in color, texture, or shape—such as sticks, bark, dried leaves, or dappled lighting patterns on vegetation—suggesting the model may rely partially on low-level visual features (e.g., texture, color patches) rather than holistic morphological recognition. In Section 5.3, we discuss approaches to reduce spurious detections

**Figure 6.**
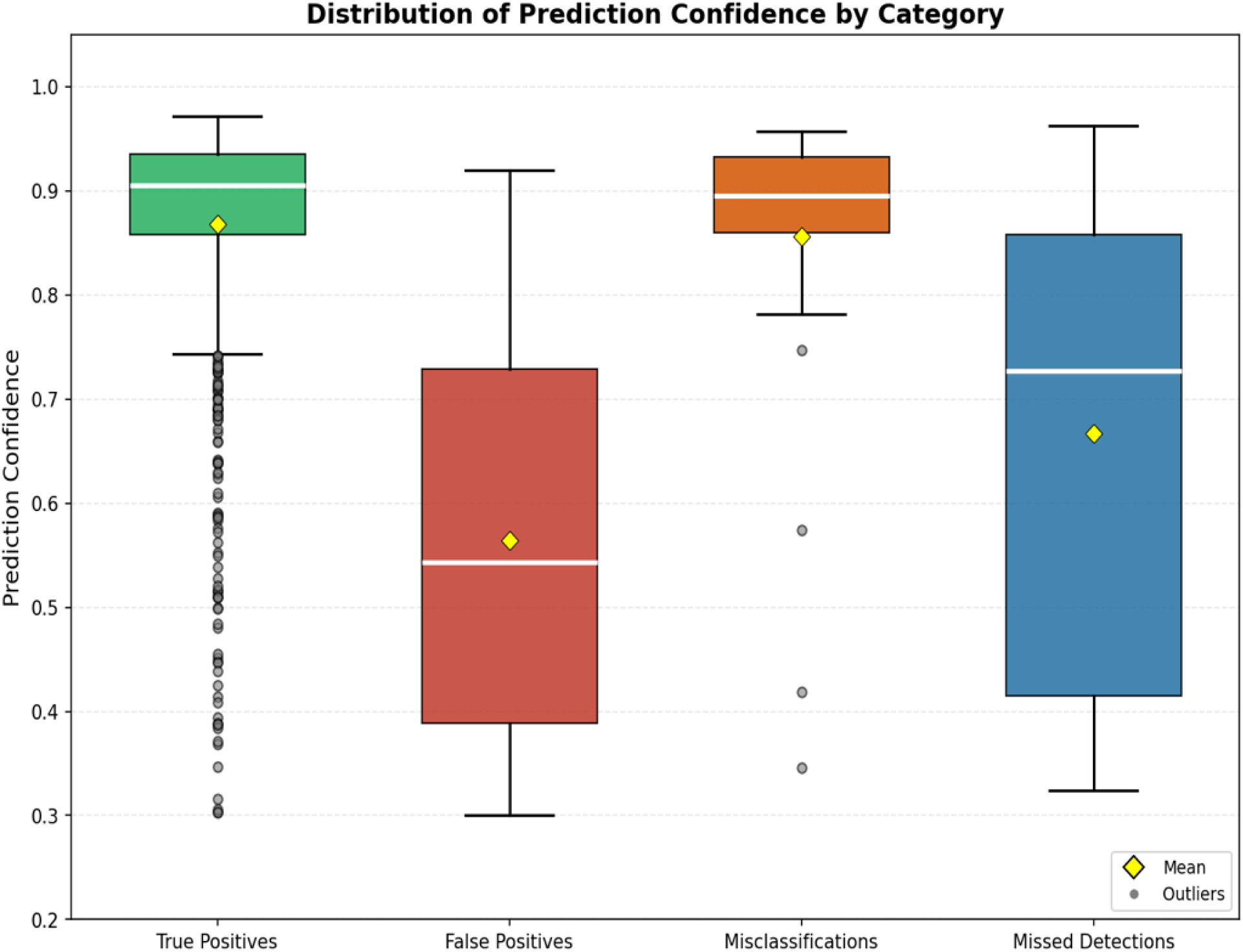
Distribution of prediction confidence across outcome categories (test set lizard instance count = 1011). Boxes span the IQR (Q1–Q3), white lines mark medians, yellow diamonds mark means, and grey dots denote outliers, defined as points falling more than 1.5 × IQR below Q1 or above Q3. True positives (n = 896, 88.6% of test set, median 0.91, IQR 0.86–0.94) and misclassifications (n = 39, 3.86% of test set, median 0.90, IQR 0.86–0.93) are tightly clustered at high confidence, whereas false positives (n = 81, 8.01% of test set, median 0.54, IQR 0.39–0.73) occupy a much broader, lower range. Missed detection statistics (median 0.73, IQR 0.42–0.86) are computed over the 44 of 67 cases for which a low-IoU prediction was produced; the remaining 23 cases are true missed detections with no prediction and thus have no confidence value.

Figure 5. Representative error modes in the detection model. (A) False-negative due to partial occlusion (n=41 images, 4.06% of test set): the model fails to detect a lizard when only a small portion of the body (e.g., head) is visible and obscured by surrounding vegetation or substrate. (B) False-negative due to multiple proximate individuals (n=9 images, 0.89% of test set): the model generates a single bounding box encompassing two closely positioned lizards rather than detecting each individual separately. (C) False-positive detection: the model incorrectly identifies non-lizard objects (e.g., sticks, bark, or lighting patterns) as lizards, often with low confidence. Blue boxes indicate predicted detections, and red boxes indicate ground truth annotations.

Figure 6. Distribution of prediction confidence across outcome categories. Boxes span the IQR (Q1–Q3), white lines mark medians, yellow diamonds mark means, and grey dots denote outliers. True positives (n = 896, median 0.91, IQR 0.86–0.94) and misclassifications (n = 39, median 0.90, IQR 0.86–0.93) are tightly clustered at high confidence, whereas false positives (n = 81, median 0.54, IQR 0.39–0.73) occupy a much broader, lower range. Missed-detection statistics (median 0.73, IQR 0.42–0.86) are computed over the 44 of 67 cases for which a low-IoU prediction was produced; the remaining 23 are true missed detections with no prediction and thus no confidence value.

#### 3.3.4. Trade-Offs and Deployment Considerations

The optimal balance between false negatives and false positives depends on the intended application context. For the Lizards on the Loose educational initiative, minimizing false negatives (ensuring detected lizards are not missed) may be prioritized to maintain student engagement and maximize capture of valid observations, even if this increases false positives that students can readily identify and dismiss during interactive review (see Section 2.6.2). Conversely, for fully automated biodiversity monitoring pipelines requiring minimal human verification, minimizing false positives should be prioritized to reduce manual review burden and prevent erroneous records from entering databases.

Our LizardLens web application currently uses confidence threshold of 0.3 and NMS IoU threshold of 0.7 values optimized for retaining true positives, but these parameters can be dynamically adjusted based on user feedback, application context, and emerging deployment priorities. The interactive review interface (Figure 2C) provides an additional quality control layer that mitigates both error types by allowing users to validate, correct, or reject automated predictions before submitting observations to online community science databases such as iNaturalist.

### 3.4. Grad-CAM: Model Interpretability

#### 3.4.1. Alignment With Known Diagnostic Features

We conducted a body-part level analysis of the Grad-CAM visualizations (n=25 images, 2.5% of the test set) to investigate the area of the image that the classification model attended to. The area of focus included the diagnostic features of the species (Table 5), as well as other body parts (e.g. front legs, hind legs, toe pads, tail) and background noises. The analysis revealed that the classification model frequently attended (n=21 images, 84% of analysed images) to morphological features known to be species-diagnostic (Table 5). However, the model did not consistently utilize all recognized diagnostic characters and, in some cases, attended to non-diagnostic features. For example, although *A. distichus* exhibits distinctive mottled body patterning (light and dark flecks), the Grad-CAM heatmap (Figure 7) attended primarily on the eyes and legs rather than body coloration. Similarly, for *A. cristatellus* (Crested anole), the model did not focus on dorso-lateral stripes that are prominent field marks in juveniles, females, and subadult males.

**Figure 7.**
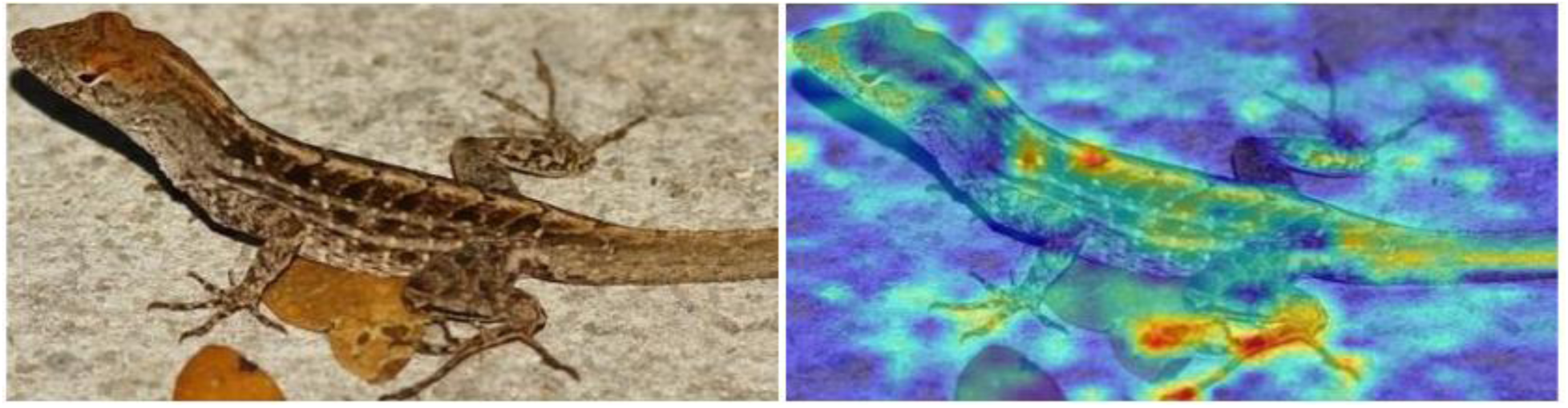
Grad-CAM visualization for a Brown Anole (Anolis sagrei). Original image (left) followed by the Grad-CAM heatmap from the Swin Transformer model (right). Warmer colors (red/yellow) indicate regions with higher contribution to the model’s prediction. The model primarily attends to the head, toepad, and limb regions rather than the dorsal patterning, indicating partial alignment with known diagnostic features.

**Table 5.** Diagnostic features used by LizardLens identify *Anolis* lizard species.

| Species | Grad-CAM diagnostic features |
| --- | --- |
| Bark anole (*A. distichus*) | Light-colored ocular ring; limb lengths |
| Brown anole (*A. sagrei*) | Variable dorsal body patterning; large hindfeet and hind toes |
| Crested anole (*A. cristatellus*) | Hindlimb length; toepad size |
| Green anole (*A. carolinensis*) | Elongated (pointed) snout; long tail; short hindlimbs |
| Knight anole (*A. equestris*) | Distinctive white shoulder stripe; pale perioral markings |

The Grad-CAM analysis provides preliminary insights into the classification model’s decision-making process but is limited by its qualitative nature and small sample size. The sample size was intentionally limited to enable careful qualitative interpretation across species, as large-scale Grad-CAM visualization is computationally straightforward but requires manual inspection to assess biological relevance. Additionally, Grad-CAM visualizes where the model focuses but does not definitively establish causal relationships between highlighted features and classification decisions (Selvaraju *et al*., 2017).

Figure 7. Grad-CAM visualizations for a Brown Anole (*Anolis sagrei*) across model stages. From left to right: original image, followed by Grad-CAM heatmaps from the four stages (Stage 1-Stage 4) of the Swin Transformer. Warmer colors (red/yellow) indicate regions with higher contribution to the model’s prediction. The model primarily attends to the head and limb regions rather than the dorsal patterning, indicating partial alignment with known diagnostic features. Original image (left) followed by Grad-CAM heatmap from the Swin Transformer (right). Warmer colors (red/yellow) indicate regions with higher contribution to the model’s prediction. The model primarily attends to the toepad and limb regions rather than the dorsal patterning, indicating partial alignment with known diagnostic features.

## 4. Discussion

Accurate species identification is a persistent bottleneck limiting the scientific utility of community science biodiversity data (Kosmala *et al*., 2016; Kelling *et al*., 2019). Here, we developed LizardLens, a two-stage machine learning pipeline that decouples object detection from species classification to identify morphologically similar *Anolis* lizard species in visually complex field photographs. Our pipeline substantially outperformed state-of-the-art single-stage architectures across all evaluation metrics, achieving 83.0% Top-1 accuracy and relative improvements of 10.5–13.2% over the best-performing single-stage model (YOLOv8x; Tables 3 and 4). The Swin Transformer classification stage achieved high precision and recall for all five species (F1 ≥ 0.85; Table 2), and Grad-CAM interpretability analysis revealed that the model independently converged on many of the same morphological features used by expert taxonomists for species discrimination (Figure 2, Table 5). Deployed as an accessible web application, LizardLens directly supports the *Lizards on the Loose* community science initiative by providing immediate, probability-based species identifications that enhance both data quality and student engagement. Below, we discuss the ecological and methodological significance of these findings and outline directions for future development.

### 4.1. Advantages of task decoupling for ecological image classification

The enhanced performance of our two-stage pipeline relative to single-stage models highlights the benefits of decoupling detection from classification in tasks requiring fine-grained visual discrimination, particularly when organisms occur in visually heterogeneous environments or when species exhibit subtle morphological differences. Prior work in fine-grained recognition has shown that accurate classification often depends on isolating discriminative regions or parts, as global image features may obscure subtle inter-class variation. In single-stage architectures such as YOLOv8x and YOLOv12x, localization and classification are learned jointly within a single model architecture, which can limit the model’s ability to focus on fine-scale features when targets are small relative to image dimensions or embedded in structurally complex backgrounds (Krause *et al*., 2014; Li *et al*., 2018; Dong, 2025). By contrast, our two-stage pipeline enables specialization of each component: the YOLOv8x detector is optimized to localize lizards independent of species identity, while the Swin Transformer classifier operates on cropped, centered representations of the target organism. This spatial decoupling reduces background interference and emphasizes diagnostic morphological features, thereby improving discrimination among visually similar species.

An additional advantage of the two-stage architecture is that it enables user interaction during the detection stage. Because localization and classification are decoupled, non-expert users can manually adjust bounding boxes even when they lack confidence in species identification. This interactive correction process allows users to improve model inputs without needing taxonomic expertise and is particularly valuable in community science and educational settings.

These advantages are particularly relevant for ecological applications, as community science photographs (e.g., iNaturalist) frequently depict small organisms in natural environments. *Anolis* lizards exemplify this challenge: they typically occupy a small proportion of a photograph and are photographed against backgrounds of bark, foliage, branches, and other structurally complex substrates that share color and textural properties with the organisms themselves, and that generate complex shadow patterns. The cropping step effectively increases the signal-to-noise ratio for the classifier by isolating the organism from its surroundings—a preprocessing strategy that mirrors the approach a field biologist would take when examining a photograph for species identification.

Notably, YOLOv12x—a more recent architecture incorporating attention-centric feature extraction designed to capture long-range spatial dependencies (Tian, Ye and Doermann, 2025)—underperformed the older YOLOv8x model across all metrics. This result suggests that architectural novelty and increased representational capacity do not necessarily translate into improved performance in ecological classification tasks that require subtle interspecific discrimination. The attention mechanisms in YOLOv12x may be better suited to tasks involving larger, more distinct objects rather than the subtle phenotypic differences characteristic of closely related species. This finding reinforces the principle that model selection for ecological applications should be guided by empirical performance on domain-specific datasets rather than general benchmark rankings (Horn *et al*., 2021; Sterkenburg and Grünwald, 2021; Tredennick *et al*., 2021).

### 4.2. Species-level variation in classification performance reflects morphological distinctiveness

LizardLens performed best for *A. distichus* (Bark anole; F1 = 0.93), a species characterized by a distinctive light-colored ocular ring and compact body proportions that distinguish it from all other Florida anoles. Similarly strong performance for *A. carolinensis* (Green anole; F1 = 0.89) and *A. equestris* (Knight anole; F1 = 0.88) likely reflects the morphological distinctiveness of these species: an elongated pointed snout and long tail in the former, and large body size with distinctive white shoulder stripes and pale perioral markings in the latter.

Conversely, the lowest performance was observed for *A. sagrei* (Brown anole; F1 = 0.85; Table 4), and the most frequent interspecific confusion (n = 16 misclassifications; Figure 4) occurred between *A. sagrei* and *A. cristatellus* (Crested anole). Rather than reflecting a simple model limitation, this confusion pattern mirrors the genuine difficulty that biology students—and occasionally lizard experts—experience when distinguishing these two species (Stroud pers. obs.). *Anolis sagrei* and *A. cristatellus* exhibit remarkable phenotypic similarity in body shape, overall coloration, and dorsal patterning, as well as semi-terrestrial habitat use (Hall *et al*., 2020; Stroud *et al*., 2020, 2024; Hall, Thawley and Stroud, 2024).

It is worth noting that *A. sagrei* also exhibits considerable sexual dimorphism in dorsal coloration and patterning, most pronounced among females (Moon and Kamath, 2019), which may further reduce classification precision by broadening the feature space the model must learn for this species. Future work could investigate whether incorporating additional information on intraspecific pattern variation as an auxiliary input improves discrimination between *A. sagrei* and *A. cristatellus*.

### 4.3. Error modes and their ecological significance

Interspecific misclassification was not the dominant source of pipeline failure (n=39 images, 3.86% of test set); rather, detection-stage errors—missed detections (n=67 images, 6.63% of test set) and spurious detections of non-lizard objects (n = 81 images, 8.01% of test set)—accounted for the majority of errors. This hierarchy of error sources carries important ecological implications. The finding that distinguishing lizards from visually similar environmental features (sticks, bark, dappled light) represents a greater challenge than discriminating among *Anolis* species, which highlights the fundamental difficulty of organism detection in structurally complex habitats—a challenge that extends well beyond *Anolis* to any small-bodied organism that relies on crypsis or occupies visually heterogeneous microhabitats (Baling *et al*., 2020; Yu, Lin and Xiao, 2024).

The two primary causes of missed detections—partial occlusion by vegetation (n=41 images, 4.06% of test set) and failure to resolve multiple proximate individuals (n=9 images, 0.89% of test set) —are both ecologically predictable. *Anolis* lizards frequently perch among branches and foliage where partial body concealment is common, and high population densities or behavioral interactions (e.g., reproductive behaviors) typically mean that conspecifics are frequently in close spatial proximity (Stroud *et al*., 2023, 2024). The Non-Maximum Suppression (NMS) post-processing step, which suppresses overlapping bounding boxes above a defined IoU threshold, appears particularly problematic when conspecifics are in close proximity. However, the practical impact of this error mode for community science applications is minimal, as most community science platforms including iNaturalist require single-species photographic records, and all multi-individual images in our dataset contained conspecifics, so missing one of two individuals of the same species does not degrade observation quality.

Spurious detections are clustered at lower confidence (n = 81, 8.01% of test set, median 0.54, IQR 0.39–0.73) as compared to confidences of true positives (n = 896, 88.6% of test set, median 0.91, IQR 0.86–0.94) (Figure 6). We use a 0.3 threshold to retain nearly all true positives, including those in the low-confidence tail, while admitting only a small residual of false positives (8.01% of the test set). These residual false positives are addressed by the interactive review interface, in which users visually inspect each bounding box and dismiss spurious detections before submission. The spuriously detected objects—sticks, bark textures, dried leaves, and dappled lighting patterns—share low-level visual features (color, texture, elongated shape) with Anolis lizards, indicating that the detection model relies partially on these coarse features rather than exclusively on holistic morphological recognition. This parallels the broader challenge of automated wildlife detection in camera trap and field photography studies, where environmental features frequently trigger false positives (Tabak, Norouzzadeh, Wolfson, Sweeney, Vercauteren, Snow, Halseth, Di Salvo, J. S. Lewis, *et al*., 2019; Maile, Duggan and Mousseau, 2023).

### 4.4. Model interpretability: convergence on biologically meaningful features

A key finding from our Grad-CAM analysis is that the Swin Transformer classification model independently learned to attend to morphological features that expert taxonomists use for *Anolis* species identification (Losos, 2009). The model attended to species-specific diagnostic features—including snout morphology, limb proportions, toepad size, ocular ring coloration, body patterning, and shoulder markings (Table 5)—that correspond closely to characters used by expert taxonomists for field identification (Krysko *et al*., 2011; Stroud *et al*., 2025). This convergence between algorithmic and expert feature use is notable because the model received no explicit guidance about which morphological characters are taxonomically informative; these features were learned entirely from the statistical structure of the training images. This result adds to a growing body of evidence that deep learning models can recover biologically meaningful signal from ecological image data (Akagi *et al*., 2020; Wei *et al*., 2022; Mostafa *et al*., 2023), and it enhances confidence that the model’s predictions are grounded in taxonomically relevant morphology rather than spurious correlations with background features or image artifacts.

### 4.5. Applications for community science and conservation

LizardLens was designed not only as a classification tool but as an integrated platform that bridges machine learning with community science education and biodiversity monitoring. The web application’s design prioritizes accessibility for non-expert users through several features: probability-ranked species predictions allow users to evaluate classification certainty, interactive bounding box correction enables users to refine automated detections, and confidence-coded color visualization provides immediate visual feedback on prediction reliability. These features are particularly important for the Lizards on the Loose initiative, where middle school students serve as primary data collectors and may lack the taxonomic expertise to independently verify species identifications (Aristeidou *et al*., 2021).

From a conservation perspective, the pipeline addresses a pressing need for high-quality occurrence data on invasive *Anolis* species in Florida. Four of the five species in our study (*A. sagrei, A. cristatellus, A. distichus*, and *A. equestris*) are non-native to Florida (Krysko *et al*., 2011; Stroud *et al*., 2019), and their ongoing range expansions have significant ecological implications for native species, including negative interactions with native *A. carolinensis* (Campbell, 2000; Stuart *et al*., 2014; Kamath and Stuart, 2015; Bush, Ellison and Simberloff, 2022). Automated tools that improve the accuracy and volume of occurrence records for these species can directly inform management strategies and contribute to invasion biology research, such as more informed invasion range forecasting (Mothes *et al*., 2019). The adjustable confidence and NMS thresholds in our pipeline allow deployment parameters to be tuned to specific use cases: lower confidence thresholds maximize detection sensitivity for engagement-focused educational settings, while higher thresholds reduce false positive rates for automated biodiversity monitoring pipelines where manual review is impractical. More broadly, by providing immediate feedback on species identifications, *LizardLens* transforms passive data collection into an interactive learning experience in which students engage directly with taxonomic decision-making—demonstrating that research-quality biodiversity data and authentic scientific experiences for young students need not be competing objectives (Shah and Martinez, 2016).

### 4.6. Limitations

Two limitations of this study should be acknowledged. First, our dataset is restricted to five *Anolis* species in Florida, and pipeline performance may not generalize to other geographic regions, taxonomic groups, or species assemblages. Second, our training data were sourced exclusively from iNaturalist “Research Grade” observations, which represent a potentially biased subsection of photographs that exclude low-quality images in which iNaturalist users were unable to collectively agree on a species’ identity. Therefore, *LizardLens* performance may degrade on images that differ substantially from those used for model training.

### 4.7. Future directions

The two-stage pipeline framework demonstrated here is readily extensible to other ecological classification challenges involving small-bodied organisms in complex habitats. For example, automated identification of frogs, insects, small birds, and small mammals from field photographs and camera trap imagery, where the same challenges of small target size, complex backgrounds, and fine-grained interspecific similarity apply (Schneider, Taylor and Kremer, 2018; Tabak, Norouzzadeh, Wolfson, Sweeney, Vercauteren, Snow, Halseth, Di Salvo, J. Lewis, *et al*., 2019; Bjerge *et al*., 2023). There are also several specific avenues for improving LizardLens. First, expanding the species set to include additional common non-native lizard species in Florida (e.g., *Agama picticauda*, *Basiliscus vittatus*, *Leiocephalus carinatus*, *Hemidactylus* sp., *Iguana iguana*, and *Ctenosaura similis*) would increase the pipeline’s utility for comprehensive biodiversity monitoring in the region (Krysko, Enge and Moler, 2019; Mothes *et al*., 2019).

Integrating geographic locality data as auxiliary model input represents a particularly promising direction for improving discrimination between morphologically similar species with partially non-overlapping ranges, such as *A. sagrei* and *A. cristatellus* (Kolbe *et al*., 2016; Stroud *et al*., 2024). Developing edge-deployed versions of the pipeline for offline use would expand accessibility to field settings with limited connectivity, while direct integration with community science platforms such as iNaturalist could streamline data submission workflows. Similarly, important insights on ecological interactions may be derived from iNaturalist images if tools are able to identify multiple species or predator-prey interactions, which are documented within our five species study system (Stroud, 2013; Ljustina and Stroud, 2016).

Finally, formal user studies with student participants in the *Lizards on the Loose* program would provide valuable data on educational outcomes, user experience, and the extent to which interactive machine learning tools enhance taxonomic learning and engagement with biodiversity science.

## 5. Conclusion

LizardLens demonstrates that decoupling object detection from species classification into a two-stage pipeline yields substantial performance improvements for fine-grained identification of morphologically similar organisms in visually complex field photographs. By achieving robust classification accuracy across five *Anolis* species while learning to attend to biologically meaningful diagnostic features, our pipeline bridges the gap between machine learning and expert taxonomic knowledge. The integration of this technical capability into an accessible, user-centered web application illustrates how computational tools can simultaneously enhance biodiversity data quality and create authentic scientific learning experiences for community science participants. As automated identification tools become increasingly central to biodiversity monitoring at scale, approaches that combine high classification performance with interpretability and accessibility will be essential for maintaining the scientific rigor and public engagement that underpin effective community science and conservation.

## Data and Code Availability Statement

The curated iNaturalist image dataset supporting this study, including bounding box annotations and train, validation and test splits, is archived in the Dryad Digital Repository (https://doi.org/10.5061/dryad.xksn02vx2).

All code for model training, evaluation, Grad-CAM analysis, and the deployed web application is archived on Zenodo (https://doi.org/10.5281/zenodo.20366694) and developed openly at https://github.com/wenhan9604/Anole_classifier. The trained model weights are included in the Zenodo archive.

## Acknowledgements

This work was supported by the David and Lucille Packard Foundation and the Maxwell/Hanrahan Foundation (both to J.T.S.). We thank Nabil Abdullah for assistance during the early development of this project. We also thank the Georgia Tech College of Computing for hosting and supporting the web application resources used in this study. This research was supported in part through research cyberinfrastructure resources and services provided by the Partnership for an Advanced Computing Environment (PACE) at the Georgia Institute of Technology, Atlanta, Georgia, USA (RRID:SCR_027619).

